# Translational predictions of phase 2a first-in-patient efficacy studies for antituberculosis drugs

**DOI:** 10.1101/2023.01.18.524608

**Authors:** Jacqueline P. Ernest, Janice Jia Ni Goh, Natasha Strydom, Qianwen Wang, Rob C. van Wijk, Nan Zhang, Amelia Deitchman, Eric Nuermberger, Rada M. Savic

## Abstract

**Background:** Phase 2a trials in tuberculosis typically use early bactericidal activity (EBA), the decline in sputum colony forming units (CFU) over 14 days, as the primary outcome for testing the efficacy of drugs as monotherapy. However, the cost of phase 2a trials can range from 7 to 19.6 million dollars on average, while more than 30% of drugs fail to progress to phase 3. Better utilizing preclinical data to predict and prioritize the most likely drugs to succeed will thus help accelerate drug development and reduce costs. We aim to predict clinical EBA using preclinical in vivo pharmacokinetic-pharmacodynamic (PKPD) data and a model-based translational pharmacology approach.

**Methods and Findings:** First, mouse PK, PD and clinical PK models were compiled. Second, mouse PKPD models were built to derive an exposure response relationship. Third, translational prediction of clinical EBA studies was performed using mouse PKPD relationships and informed by clinical PK models and species-specific protein binding. Presence or absence of clinical efficacy was accurately predicted from the mouse model. Predicted daily decreases of CFU in the first 2 days of treatment and between day 2 and day 14 were consistent with clinical observations.

**Conclusion:** This platform provides an innovative solution to inform or even replace phase 2a EBA trials, to bridge the gap between mouse efficacy studies and phase 2b and phase 3 trials, and to substantially accelerate drug development.

## Introduction

*Mycobacterium tuberculosis* remains one of the deadliest infectious agents globally. Tuberculosis (TB) drug discovery and development activity has increased emphasis on shorter, more universal regimens to treat all TB cases independent of resistance status^1,2^. However, with an increasing number of new drugs and limited resources for clinical trials, further innovation of drug development is imperative to identify effective drugs and regimens more efficiently and with higher confidence^1–3^. A phase 2a early bactericidal activity (EBA) study is typically the first clinical evaluation of novel anti-TB drug efficacy with the primary purpose of detecting the presence and magnitude of EBA and informing possible dose-response relationships^4^. However, the cost of phase 2a trials can range from 7 to 19.6 million dollars on average, while more than 30% of drugs fail to progress to phase 3^5^. This highlights the challenges inherent in translating results in preclinical models into successful clinical outcomes. Traditional translation of findings from preclinical *in vivo* models, by pharmacokinetic modeling and allometric scaling to identify the dosing regimen in humans that best matches the efficacious drug exposure in animals, is insufficient. Mechanistic mouse-to-human pharmacokinetic-pharmacodynamic (PKPD) models that describe the bacterial kill and PKPD relationships are better at predicting clinical results, including the results of late-stage trials^6–8^. Therefore, our objective is to establish a relevant and robust model-based translational platform that can reliably link preclinical to clinical drug development and predict early efficacy trials for anti-TB drugs across different compound classes (Figure 1). We compiled a comprehensive preclinical and clinical database of PK, PD, and baseline bacterial growth data for nine drugs. The drugs used to develop and validate our proposed platform were rifampin (RIF), isoniazid (INH), pyrazinamide (PZA), rifapentine (RPT), bedaquiline (BDQ), delamanid (DLM), pretomanid (PMD), moxifloxacin (MXF) and linezolid (LZD).

**Figure 1.**
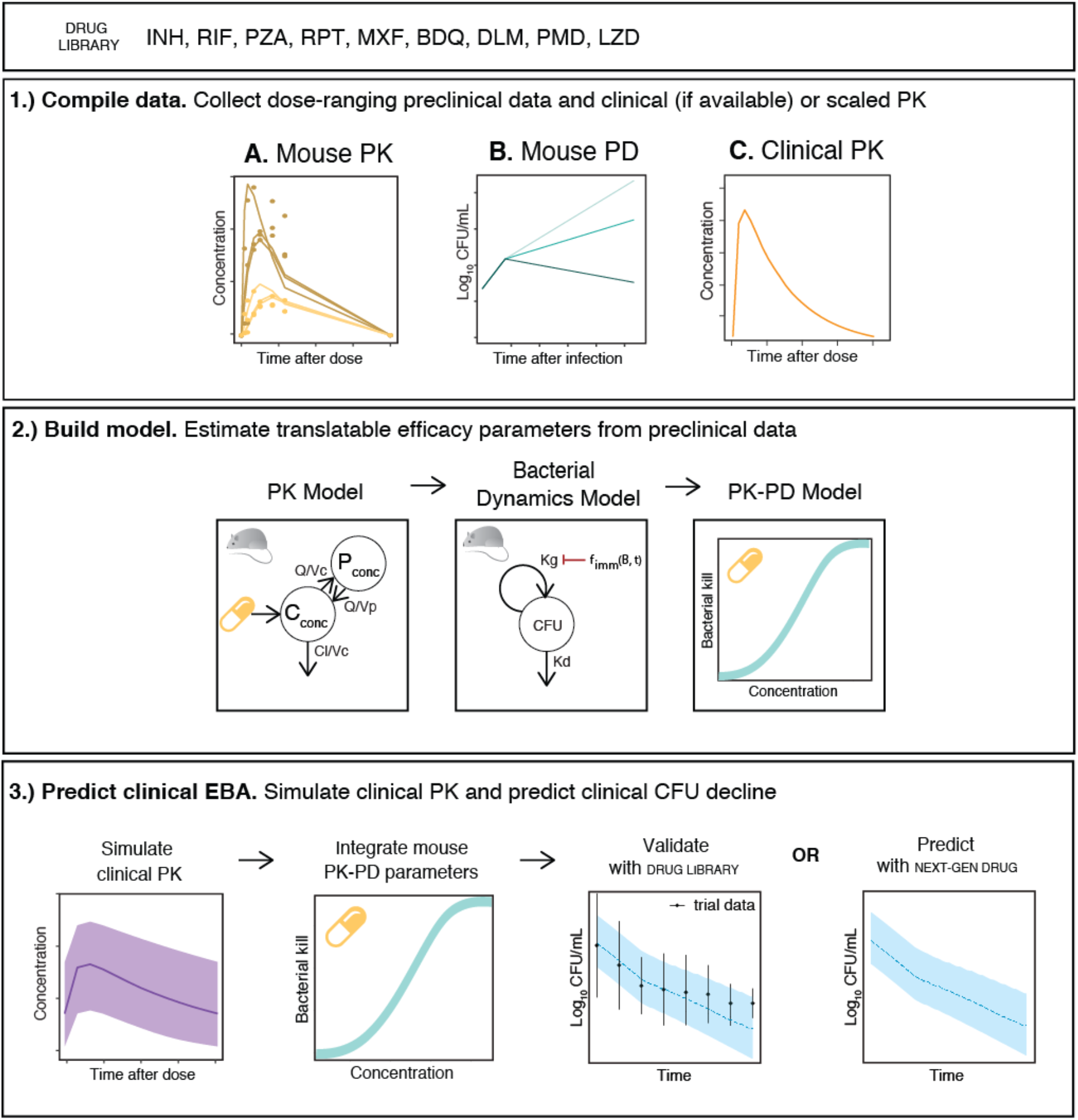
The translational pharmacology approach to predicting early bactericidal activity in patients. Components necessary for translation include mouse PKPD and clinical PK (actual or scaled). The estimated relationship between drug concentration and bacterial kill is assumed to be portable after correction for protein binding and integrated with clinical PK. Using baseline bacterial burden from previous EBA trials as initial conditions, the early bactericidal activity is simulated with the translational model.

The translational platform in the present study intends to increase the accuracy of preclinical to clinical translation by enabling quantitative prediction of clinical studies from preclinical outputs and serves as a foundation for model-informed TB drug discovery and development.

## Methods

### Drug dataset for model building and validation

To build our model and evaluate its predictive accuracy for clinical EBA, nine first- and second-line anti-TB drugs (BDQ, DLM, INH, LZD, MXF, PMD, PZA, RIF, RPT) were selected for which mouse PK, mouse PD, human population PK models and human clinical EBA data were available.

### Data required to assess preclinical drug efficacy

A large database of PK and PD data in mice was collected (Figure 2, Table S1). PK experiments in BALB/c mice were dose-ranging (2-10 dose levels), single or multiple oral dosing for up to 8 weeks, with 29-238 observations of plasma concentration per drug. PD experiments in BALB/c mice infected through aerosol delivery were dose-ranging (2-15 dose levels) with treatment durations of 21-70 days, and 55-252 observations of lung CFU counts per drug. Most experiments were performed at Johns Hopkins University (Table S1). Lung CFU counts were measured by plating lung homogenates at designated time points. In case of murine data showing unexpected trends such as a double peak per oral dose in a PK profile e.g. DLM mouse PK (Figure 2a), data accuracy was confirmed in collaboration with experimentalists.

**Figure 2.**
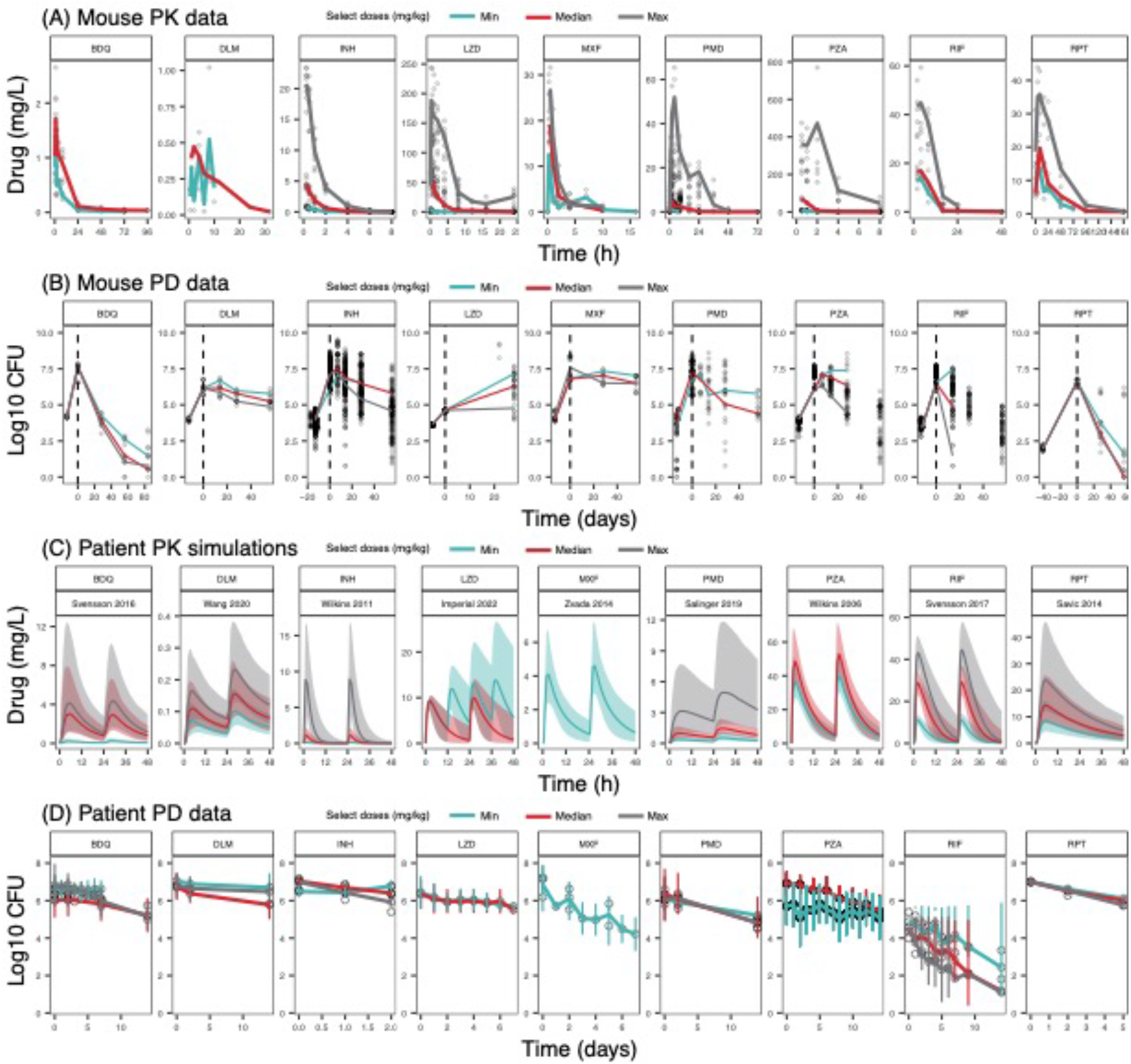
A rich dataset of mouse and human PK and PD data for 9 first- and second-line TB drugs was compiled for model building. Only minimum, median and maximum doses are represented as median lines when multiple doses were present. Data points for all doses are plotted. Information on all doses is present in Table 1. (A) Mouse pharmacokinetic (PK) data presented for the following doses: BDQ 12.5, 25 mg/kg; DLM 2.5, 3 mg/kg; INH 1.56, 6.25, 25 mg/kg; LZD 5, 100, 500 mg/kg; MXF 100, 200, 400 mg/kg, PMD 6, 28.8, 486 mg/kg; PZA 7, 100, 900 mg/kg; RIF 10, 15, 40 mg/kg; RPT 5, 10, 20 mg/kg. All doses were given once daily unless otherwise stated. (B) Mouse pharmacodynamic (PD) data presented for the following doses: BDQ 12.5, 25, 50 mg/kg; DLM 3, 10, 100 mg/kg; INH 0.1, 6.25, 100 mg/kg; LZD 100, 300, 1000 mg/kg; MXF 25, 50, 100 mg/kg; PMD 6.25, 30, 600 mg/kg; PZA 3, 50, 900 mg/kg; RIF 2.5, 40, 640 mg/kg; RPT 5, 10, 20 mg/kg. All doses were given once daily, 5 days a week, unless otherwise stated. (C) Human PK simulations from validated population PK models presented for the following doses: BDQ 25, 200, 400 mg; DLM 100, 200, 400 mg; INH 9, 75, 600 mg; LZD 600 mg once daily, 600 mg twice daily; MXF 400 mg; PMD 50, 200, 1200 mg; PZA 2000 mg; RIF 600, 1350, 1950 mg; RPT 300, 600, 1200 mg. All doses were given once daily, unless otherwise stated. (D) Human Phase 2a early bactericidal activity study data presented for the following doses: BDQ 25, 200, 400 mg; DLM 100, 200, 400 mg; INH 9, 75, 600 mg; LZD 600 mg once daily, 600 mg twice daily; MXF 400 mg; PMD 50, 200, 1200 mg; PZA 200 mg; RIF 600, 1350, 1950 mg; RPT 300, 600, 900, 1200 mg. All doses were given once daily, unless otherwise stated.

**Table 1.**
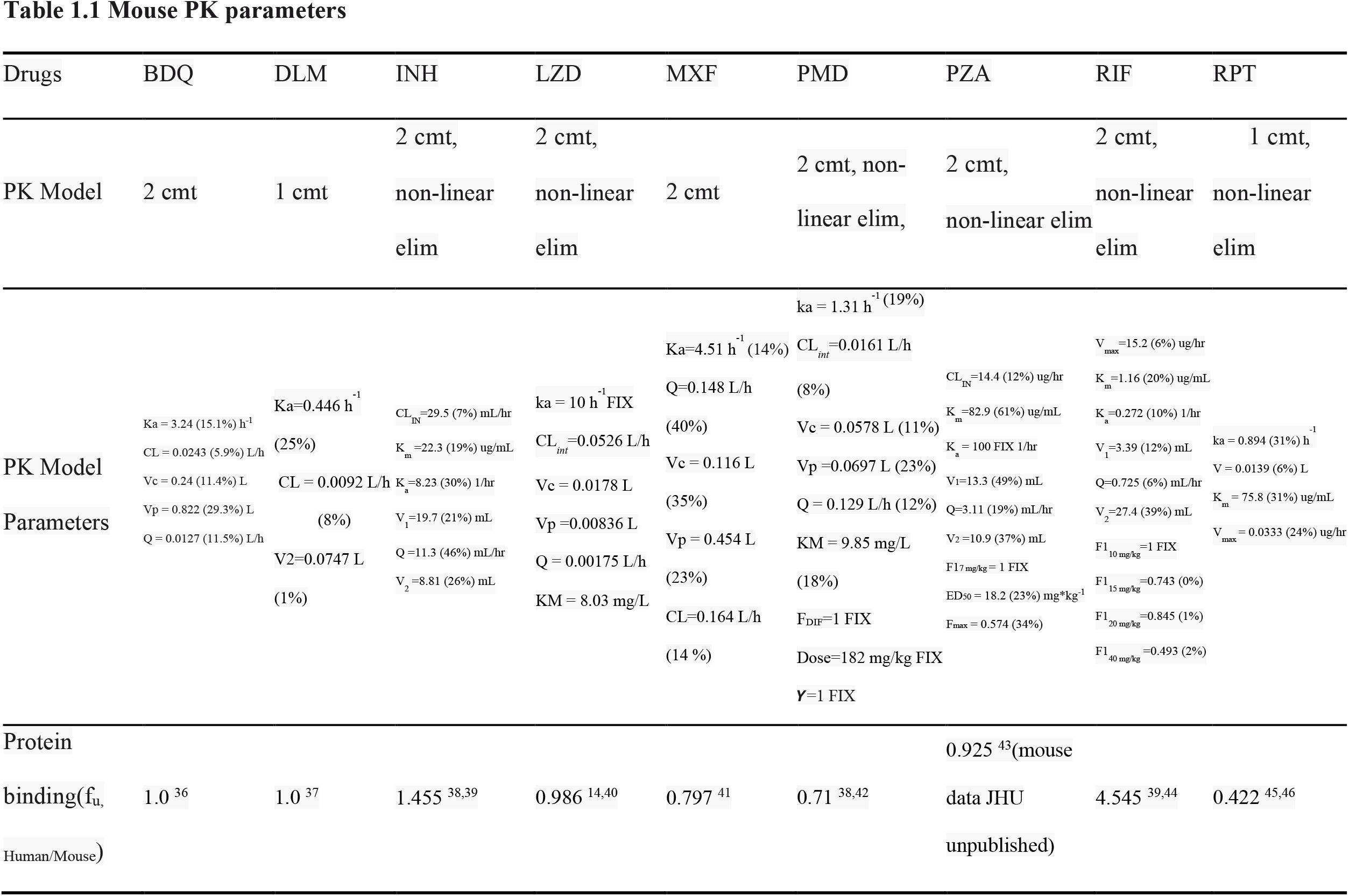
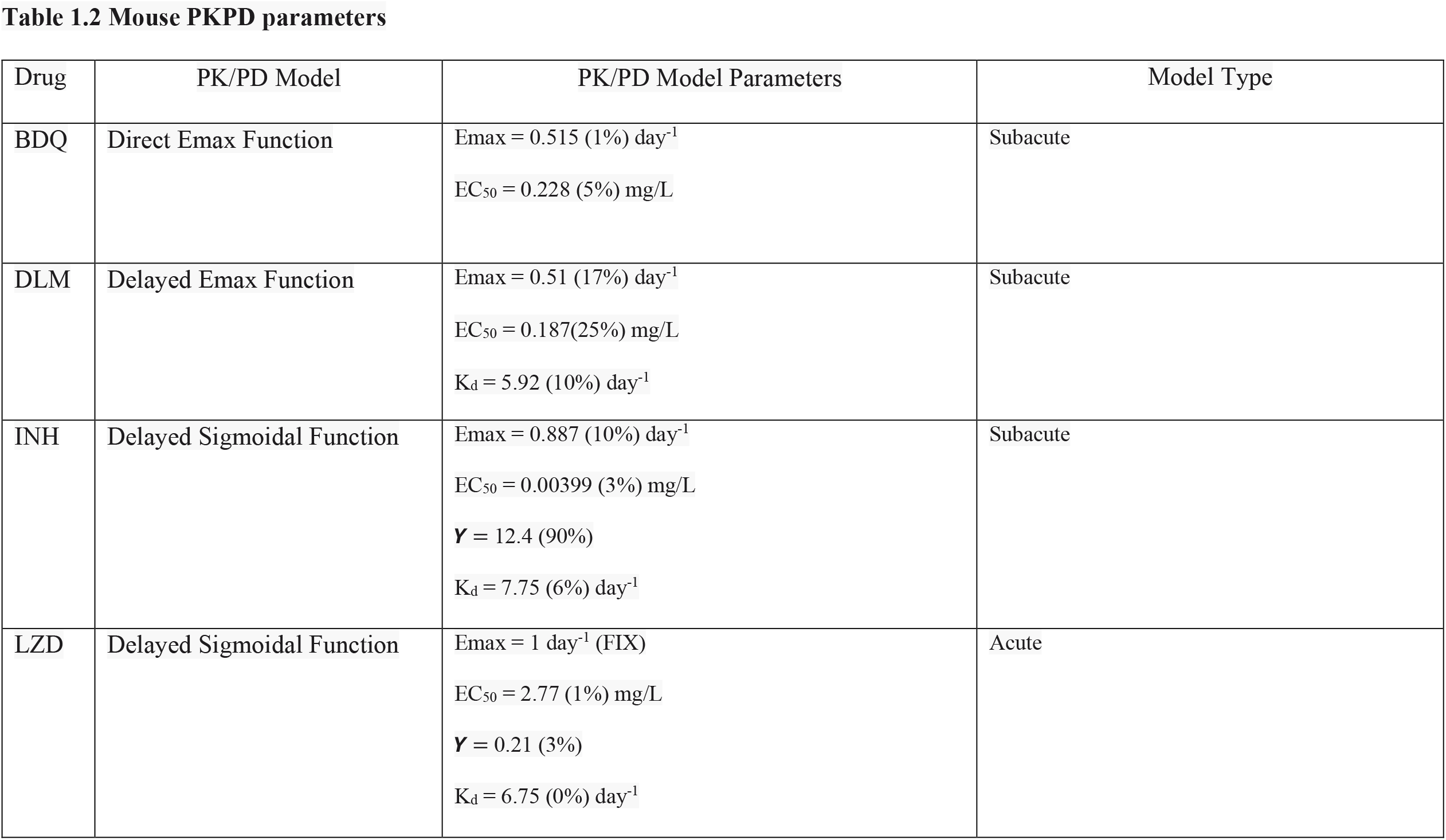
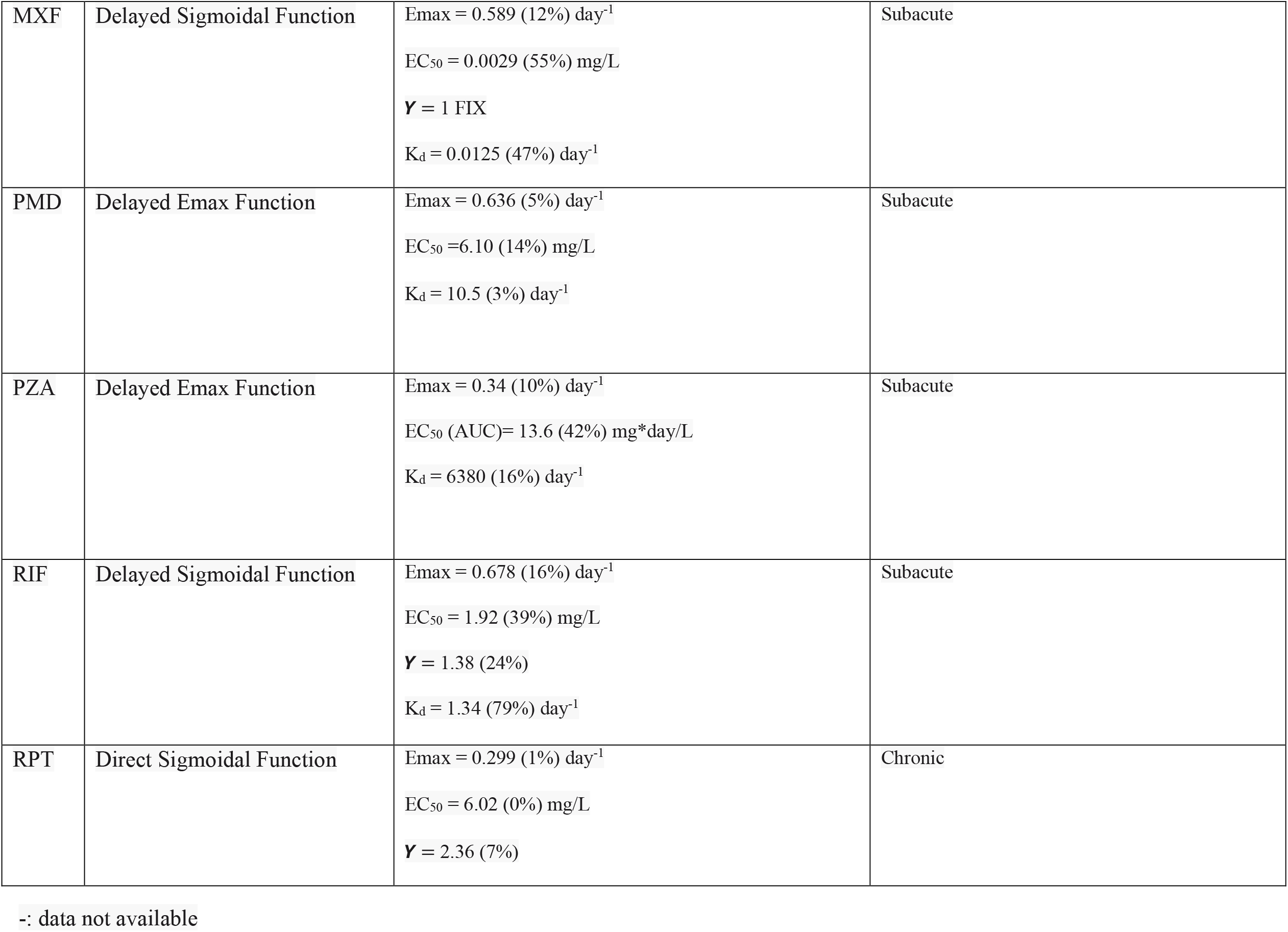
Parameter estimates of final PK and PKPD models for nine TB drugs in mouse studies.

### Mouse PKPD model development

An integrated mouse PKPD model was developed for each drug. PK data were described using one- or two-compartment models with first order absorption with or without delay, and saturable elimination when necessary. The bacterial growth dynamics without treatment was described using our previously published baseline model (Eq. S1)^9^. The baseline model captures the decreased rate of growth over time and attributes the decline to time- and bacteria-dependent immune control over the infection. The drug effect, measured as the log_10_ CFU drop independent of the immune effect over time, was incorporated using a sigmoidal Emax relationship (Eq. S2). A delay effect (Kd) was included to mouse PKPD models to establish an indirect relationship between plasma drug concentrations and drug effect at the site of action (Eq. S3 & S4). Detailed model development and model diagnostics can be found in supplemental materials.

### Prediction of the outcomes for clinical EBA studies

The PKPD relationship quantified in mice was used to predict the clinical EBA. Drug concentrations in humans were simulated based on clinical population pharmacokinetic models (Table S1) to drive the concentration-effect relationship in the clinical predictions. Where clinical population PK models were unavailable, allometric scaling from mouse PK was used^10^. Protein binding ratios between humans and mice 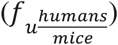 were used to convert unbound plasma drug concentrations from human to mouse to translate the mouse PKPD relationships (Table S1)^11–17^.

Clinical predictions for 9 drugs were simulated, with 14 unique studies at several dose levels were used for validation. Predictions were done by simulating CFU decline in 1000 virtual patients treated with the same dose as reported in the clinical EBA study. The baseline (Day 0) sputum values used were derived from the mean value for each arm reported in each study, and the variability in baseline bacterial burden between individuals used was the baseline variance among all clinical studies. The net growth and death of bacteria without treatment was assumed to be zero (Eq. S5). Predictions were reported as the mean and standard deviation of the predicted time course of CFU decline. For drugs where observed data were available, the data were overlayed for visual inspection. Finally, quantitative predictions of commonly reported parameters (change from baseline to Day 2 and from Day 2 to Day 14) were compared to the observed at various dose levels along a line of unity.

### Software and Statistical method

Preclinical and clinical PKPD modelling was performed in NONMEM (7.4.3) through PsN (4.8.1.). For LZD preclinical PK, Monolix (5.0.0) was used. Models were developed following numerical and graphical diagnostics, assessing drop in objective function value through the likelihood ratio test and parameter precision, as well as goodness-of-fit plots and visual predictive checks, respectively, in addition to pharmacological relevance. Data transformation and graphical output were performed in R (4.1.3) through the RStudio (2022.02.3) interface using the xpose4 and tidyverse packages.

## Results

### Large preclinical and clinical PK and PD database of nine TB drugs

We collated a rich longitudinal dataset of mouse PK (plasma concentrations, 1146 data points) and PD data (lung CFU counts, 4042 data points), as well as human population PK models and human PD data (sputum CFU counts) (Table S1). PD experiments were done mostly in mouse infection models infected via aerosol with an inoculum size no less than 3.5 log_10_ CFU/ml and incubation periods of 13-17 days, prior to the start of treatment. Exceptions were LZD, which had an incubation period of 5 days, but had a similar inoculation size of larger than 3.5 log_10_ CFU/ml, and RPT which had an incubation period of 41 days but a lower inoculation size than 3.5 log_10_ CFU/ml.

Human PK data were simulated using published models from literature (Table 1 and Figure 2C). Human PD data with a total of 260 human sputum CFU datapoints originating from Phase 2a trials across 13 different studies ranging from 2 to 14 days were used to validate our Phase 2a EBA predictions.

### Preclinical PK and PKPD models adequately described mouse data

The final PK and PKPD model parameter estimates are shown in Table 1. A 2-compartment model with saturated clearance described via the Michaelis Menten equation best described the mouse plasma concentration data for BDQ, INH, LZD, PMD, PZA and RIF. MXF was best described using a 2-compartment model with linear elimination, RPT by a 1-compartment model with saturated elimination, and DLM by a 1-compartment model with linear elimination. Visual predictive checks of the final model for both mouse PK and PKPD data showed good fits (Figures S1 & S2). The exposure-response relationships for each drug in mouse infection models are summarized in Table 1 and Figure S4 and aligned with clinical knowledge of the efficacy of each drug.

### Clinical EBA was well predicted by translational platform

The translational platform predicted clinical EBA in TB patients receiving monotherapy with the nine drugs as shown in Figure 3. Our predictions overlapped well with the observed data across multiple doses and timepoints for most of the drugs. BDQ and LZD had slight overpredictions at the later time, and RPT showed activity up to 5 days after a single dose, whereas our model predicted limited declines in CFU.

**Figure 3.**
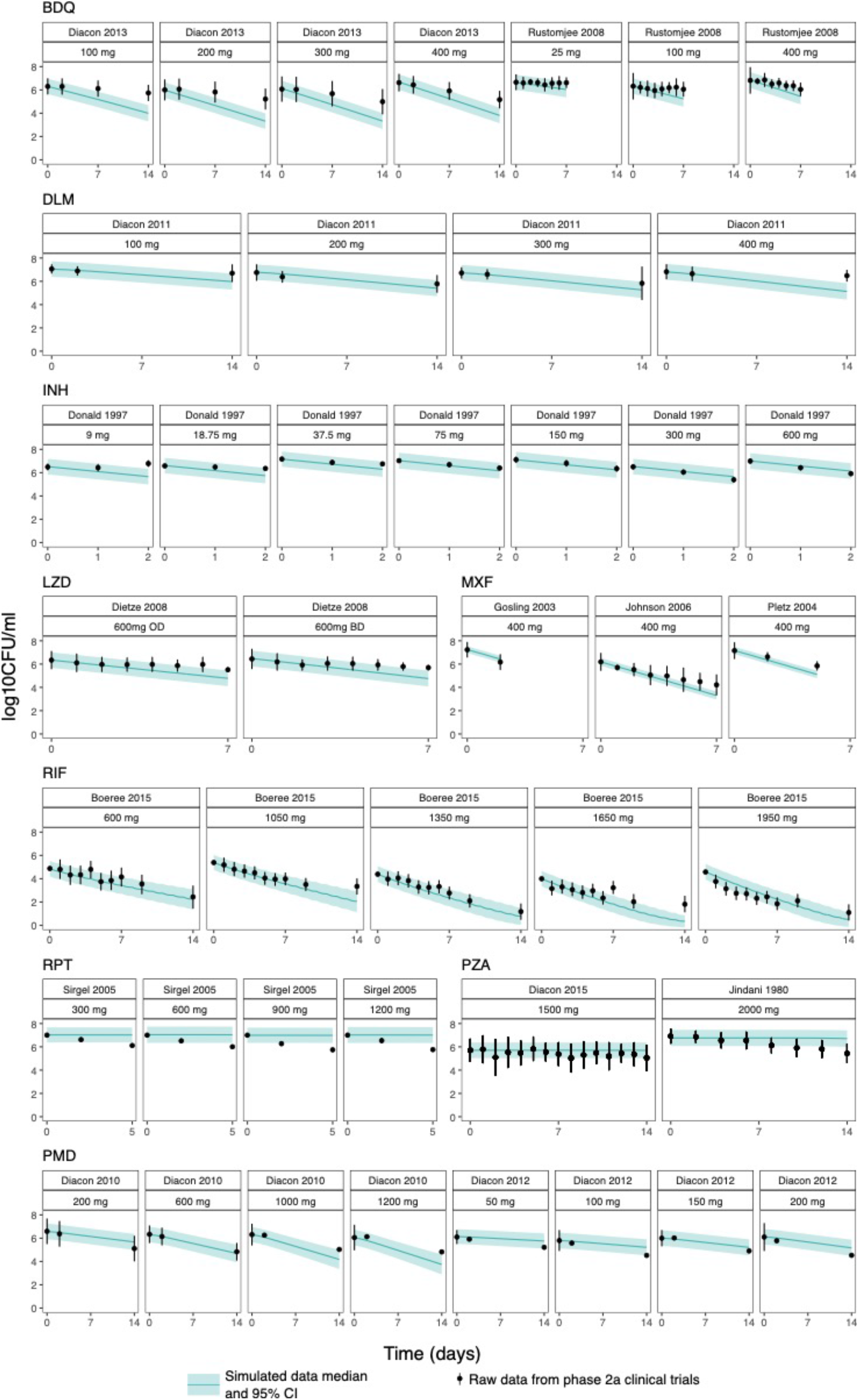
Translational (mouse to human) PKPD model predicts clinical EBA trial results well. Medians and 95% confidence intervals of 1000 simulations from the translational model overlap with observed EBA data from clinical trials.

Agreement between predicted and observed quantitative change in CFU is shown in Figure 4 as a correlation plot for EBA at time intervals of 0-2 days and 2-14 days. Most predictions for 0-2 days fell within 0.25 log_10_ CFU/ml/day of the observed EBA as indicated by the line of unity and corresponding dotted lines. Predictions for 2-14 days were even closer to observed. Predictions were overall consistent with the observed data in the clinical EBA studies for all nine drugs, except for RPT where activity was underpredicted.

**Figure 4.**
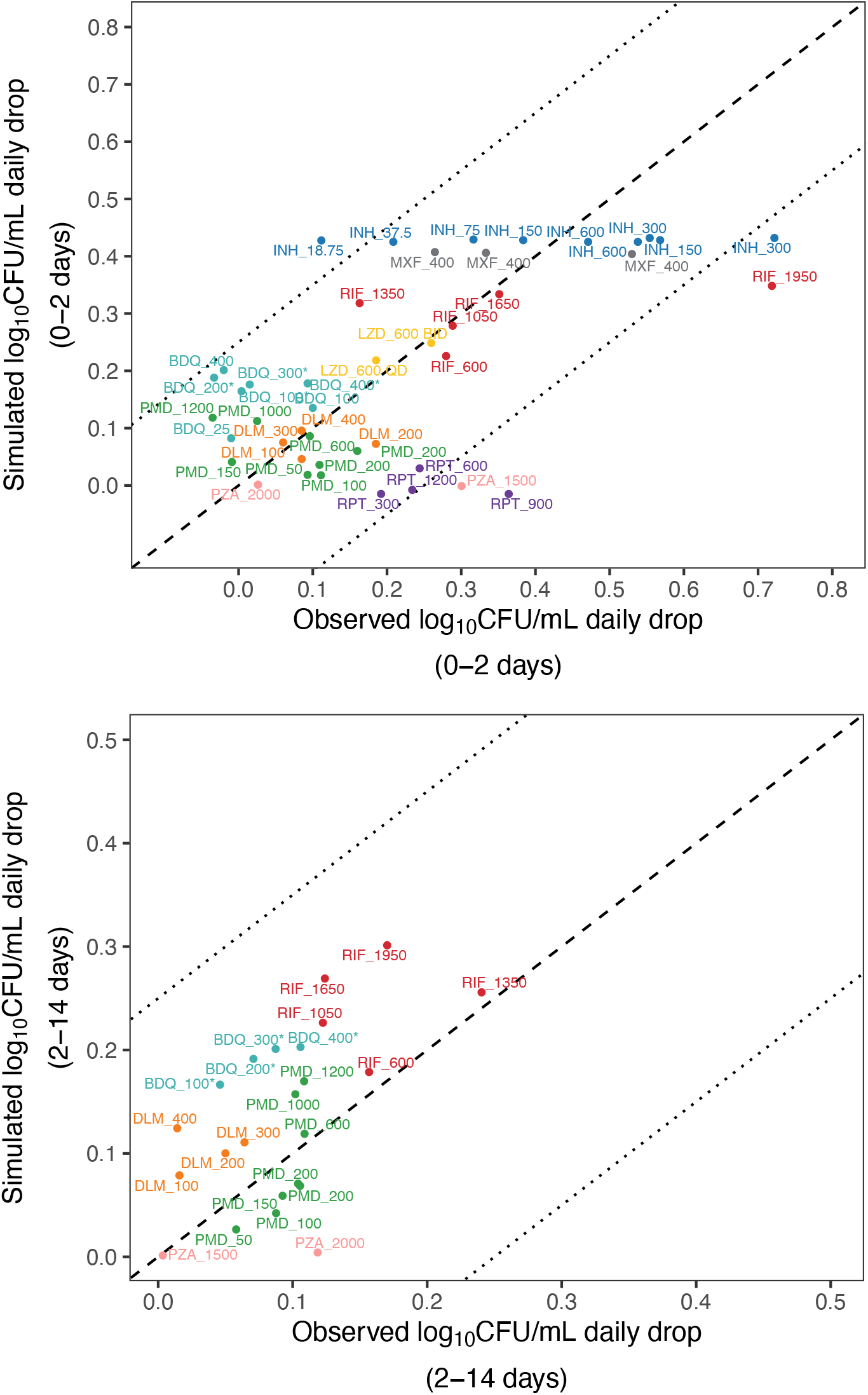
**Model-based prediction of daily change in log**_**10**_ **CFU/mL correlates well with clinically observed daily change in log**_**10**_ **CFU/mL for nine TB drugs at multiple dose levels of monotherapy between Day 0 to 2 (top) and Day 2 to 14 (bottom)**. For some drugs, Day 14 data were not available. Line of unity (dashed line) ± 0.25 (dotted lines). BDQ = bedaquiline, DLM = delamanid, INH = isoniazid, LZD = linezolid, MXF = moxifloxacin, PMD = pretomanid, PZA = pyrazinamide, RIF = rifampin, RPT = rifapentine. *regimen contained a loading dose

## Discussion

We established a mouse-to-human translational platform by integrating a bacterial dynamics model, mouse PKPD relationships, clinical PK and species-specific drug plasma protein binding and validated the platform with clinical EBA data (Figure 1). The changes in sputum CFU counts over the first two days and from Day 2 to Day 14 in TB patients receiving monotherapy with each of nine TB drugs in 13 clinical EBA studies were successfully predicted, except for RPT (Figure 3 and 4). Of the clinical EBA studies included in our analysis, the RPT EBA trial was the only one in which EBA was assessed for multiple days after a single dose. Our human population PK model indicated RPT was mostly cleared from the body two days after a single dose, but the trial results indicated RPT was still exerting an effect on bacterial load between two and five days post-dose. It is possible that RPT has a post-antibiotic effect that was not sufficiently captured by the model^18^. The model overpredicted the EBA of BDQ. However, in the model, the active metabolite, BDQ-M2, was not considered. In mice, M2 is estimated to contribute approximately 50 percent of the drug effect. One possible reason for the overprediction are the parent-to-metabolite ratios between species differ. Future studies can account for these differences.

Murine TB models are routinely and often exclusively used as *in vivo* efficacy models in preclinical TB drug development^19^. As the inoculum size and incubation period for bacterial infection in the lung prior to treatment can affect drug response^9^, we standardized our inclusion criteria to experiments using the most common design with the incubation duration of 13-17 days and inoculum size to larger than 3.5 log_10_ CFU/ml. Incubation durations outside this range were considered only when data were otherwise not available, which was the case for LZD and RPT.

A key component to our model accuracy is the addition of the bacterial dynamics model. Mouse and human immune activation against TB infection differ significantly, therefore the underlying baseline of bacterial dynamics will differ. Subtracting the mouse immune effect on bacterial decline more accurately estimates the drug contribution to CFU decline. Without such consideration, the clinical CFU decline is overpredicted (Figure S3). Despite inherent differences between species in terms of drug PK, sampling (whole lung homogenate versus sputum), and infecting bacterial strain, the relationship between drug effect on bacteria and the concentration to achieve the effect appear, based on this analysis, to be portable between mice and patients. In addition, although the mouse strain used in the studies (BALB/c) models intracellular bacteria but not extracellular bacteria in caseous lesions^20^, the PKPD relationships observed in this model, when derived in comparison to the baseline bacterial dynamics, appear to accurately reflect those observed in EBA studies. Other approaches or more information may be needed to fully account for drug exposures at the site of infection in cavities or other caseous lesions or any PK/PD relationships unique to those microenvironments. ^3,21,22^

Clinical EBA was predicted well across 14 studies spanning more than two decades. Compared to the participants enrolled in more recent EBA studies (2007 to 2015)^23–29^ at the same site, the participants enrolled between 1992 and 2005^30–34^ had more severe disease and therefore higher baseline CFU counts in their sputum samples (mean baseline: 6.9 log_10_ CFU per mL). However, the predictive accuracy of our model was robust despite this large variation in baseline bacterial burden. For example, RIF had a good overlap of predicted and observed EBA (Figure 3) despite the study being conducted in 2015 with the lowest median baseline of 4.58 log_10_ CFU per mL^24^.

Clinical EBA studies are the only acceptable way to evaluate a drug as monotherapy in TB patients despite their limitations on predicting long-term efficacy. In addition to detecting the presence of an EBA response, the trial can inform on the dose-response curve (e.g., INH and RIF), which could be used in dose selection for future trials^22,24,35^. We have shown here that our translational platform can adequately predict these outcomes. With limited resources, this costly clinical study can be designed more efficiently or avoided altogether by using our approach to predict a reliable result regarding clinical dose-response effects, and to provide useful information about dose and/or drug candidate selection for further clinical development. This scenario is well exemplified by the nitroimidazole, PMD. PMD has a dose response at doses up to 192 mg/kg in mice which, following the conventional allometric scaling method, approximates 1500 mg in humans. However, such translation is problematic as the clinical observations from two human EBA trials demonstrated no dose response above 200 mg in human EBA. Using our translational platform, we found that the drug effect of PMD reaches plateau after 200 mg which is consistent with clinical observations (Figure S4). Therefore, our translational platform could serve as a powerful tool for, but not limited to, better dose selection for clinical trials design. By better informing dose selection, the translational modeling platform may reduce the time and effort spent in early clinical development, and therefore, accelerate progress to trials that are more informative of long-term outcomes.

Building on our translational framework, efficacy of combination regimens of TB drugs tested preclinically can be predicted in future work. This shows the principles of how preclinical data used in a model-based translational framework can inform the design of clinical late-stage efficacy studies, such as phase 2b studies. Future goals to improve the platform include characterizing PKPD relationships of combination regimens by accounting for PKPD drug-drug interactions, as well as characterizing lesion-specific PKPD relationships. Clinical TB disease (e.g., caseation necrosis and cavitation) will be represented in the translational platform to include infection and efficacy data in animal models of TB with more human-like necrotic lesions, such as C3HeB/FeJ mice and New Zealand white rabbits^21^. Our translational platform may then be able to predict late-stage trials of combination regimens. If so, our platform could reduce dependence on phase 2a efficacy studies by predicting EBA and also directly inform the design of phase 2b and phase 3 studies to assist clinical anti-TB regimen development.

In summary, we established a foundation for translating the results from mouse efficacy models to clinical EBA studies through establishing quantitative relationships involving mouse PK and PD, as well as drug dose response *in vivo*. In the future, our platform will be expanded to include combination regimens and longer durations of treatment by accounting for PKPD drug-drug interactions, and necrotic lesion penetration. This platform is an innovation to accelerate TB drug development and a good example of model-informed drug discovery and development.

## Acknowledgements

We acknowledge Kelly Dooley for insightful discussions, and TB Alliance for support and generously sharing the in-house data of anti-TB drugs linezolid and pretomanid. This work was supported by NIH Grant R01 AI-111992.

## Author contributions

The manuscript was written by JE, JG, NS, QW, RW, and NZ and commented on by all authors. JE, JG, NS, QW, RW, and NZ contributed to data collection, model development, data and model management, and code review. Contributions to data collection and model development were as follows, JE: BDQ and RPT; NS: LZD; QW: DLM, PMD, and MXF; NZ: PZA, RIF, and INH. JG and RW revised the manuscript, including generation of figures and tables, and carried out code review for all drugs. AD worked on data collection, human PK model development, simulation, and preliminary model development. EN provided preclinical data used in our current study, provided substantial scientific context, and edited the manuscript. RS supervised the whole research.

## Supporting information

### Supplemental Methods

**Figure S1 Visual predictive checks for final mouse PK models at representative doses**

**Figure S2 Visual predictive checks for final mouse PD models at representative doses**

**Figure S3 Comparison between human plasma drug concentrations reached at clinical dose levels (light grey), upper limits of drug concentrations within safety ranges (dark grey) and concentration-response relationships for nine TB drugs**

**Figure S4 The immune component of the model-based translational platform is essential for accurate prediction of early bactericidal activity**

**Table S1 Mouse and human PK and PD database of nine TB drugs**

